# ICE BINDING PROTEIN ACTIVITY IS COMMON AMONG TEMPERATE PACIFIC INTERTIDAL INVERTEBRATES

**DOI:** 10.1101/2025.04.08.647825

**Authors:** N.H.W. Moyes, A.K. Bertram, K.E. Marshall

## Abstract

Intertidal invertebrates often adopt freeze tolerance as a strategy to survive winter low tides, but the physiological mechanisms by which intertidal species survive freezing are not well understood. Ice binding proteins (IBPs) may play an important role, but their occurrence throughout the intertidal zone worldwide is not well-catalogued. Here we survey species in the intertidal zone of Vancouver, BC to <u>assess IBP activities</u> and the <u>possible role they play in freeze</u> <u>tolerance</u>. We conducted targeted assays in multiple intertidal species to measure three distinct activities of IBPs: ice nucleation, ice recrystallization inhibition, and thermal hysteresis. We also exposed intertidal species to a cold exposure mimicking a freezing event in the intertidal and <u>measured survival post-exposure</u>. We found that IBP activity was widespread across intertidal species, and that species with the highest degree of freeze tolerance also display ice nucleating activity attributable to proteins. We also found that species that live in the high intertidal appear to show more IBP activity than species that live lower in the intertidal. This work significantly increases the number of known species containing IBPs and provides a framework for sampling new species for IBP activity in the intertidal and elsewhere.

## Introduction

Intertidal species live between the boundaries of high and low tides, cycling between aerial emersion and saltwater immersion. Many intertidal invertebrates have very low mobility, or are entirely sessile, making behavioural thermoregulation to avoid temperature extremes very difficult. In temperate and polar regions of the world, these animals can be exposed to sub-zero air temperatures during the winter months. While there is long-standing interest in the evolution of stress tolerance in this unique habitat, how these animals survive the cold is poorly understood (Gill et al., 2024).

Cold-hardy species across habitats largely use one of two strategies to survive the cold: avoid freezing through a suite of physiological or behavioural mechanisms (freeze avoidance), or control internal ice growth and make freezing survivable (freeze tolerance). Freeze avoidance is often achieved via colligative freezing point depression and behavioural avoidance of ice (Lee, 2010). Due to the limited mobility of many intertidal invertebrates, constant presence of external ice nucleators, and the fact that they are osmoconformers, achieving freeze avoidance is challenging if not impossible. Thus, many intertidal invertebrate species are freeze tolerant (Gill et al., 2024; Storey and Storey, 2013). Although freeze tolerance is common among temperate and polar intertidal invertebrates, the physiological and biochemical mechanisms they use to achieve freeze tolerance are not well-understood and represent a unique evolutionary path to achieving this physiological feat (Aarset, 1982; Gill et al., 2024).

One mechanism for achieving freeze tolerance is the expression of high molecular weight cryoprotectants such as ice binding proteins (IBPs). IBPs function by binding to the surface of an ice crystal and manipulating its growth (Bar Dolev et al., 2016). There are three main classes of IBPs based on their activity: thermal hysteresis (TH) proteins, ice recrystallization inhibition (IRI) proteins, and ice nucleating proteins (INPs). TH proteins create a gap between the freezing and melting points of ice, making an energetic barrier to growth of ice crystals (Kristiansen and Zachariassen, 2005). These proteins are primarily found in species that avoid freezing, since their primary role is to prevent ice formation. IRI proteins act to prevent the recrystallization of small ice crystals into thermodynamically-favoured large crystals (Capicciotti et al., 2013). These proteins are used primarily to prevent mechanical damage from large and sharp ice crystals, and are thought to play an important role in freeze tolerance (Knight and Duman, 1986). Finally, ice nucleation proteins induce ice growth at high sub-zero temperatures and are used by freeze-tolerant species to control when and where ice grows inside their body (Bar Dolev et al., 2016). INPs can be used to keep ice growth extracellular, as intracellular ice is fatal in most animals (Sinclair and Renault, 2010). IBPs have evolved independently multiple times across all kingdoms of life and are a well-established way for freeze-avoidant and freeze-tolerant species to survive the cold (Bar Dolev et al., 2016; Vance et al., 2019).

Despite the prevalence of IBPs across taxa, there are only six documented cases in intertidal species (Gill et al., 2024). Despite the rarity of documented IBP activity in intertidal species, there is genomic evidence that IBPs are much more widespread across the intertidal environment, and that IBPs are more likely to evolve in the intertidal that in other habitats (Box et al., 2022). Four of the documented IBPs in intertidal invertebrates come from the Atlantic coast, and are all INPs found in Molluscs (Aunaas, 1982a; Aunaas, 1982b; Loomis and Zinser, 2001; Madison et al., 1991). There is only one known example of a TH protein in an intertidal species, found in an Antarctic nemertean (Waller et al., 2006). *Mytilus trossulus* is the only

Pacific invertebrate with a confirmed IBP (Box, 2021). There is some molecular evidence that the Manila clam *Ruditipes philippinarum* possesses an IBP, but its activity has not been measured (Dong et al., 2022). Given the high likelihood of IBPs occurring in the intertidal zone, it is plausible that many freeze-tolerant animals in the intertidal have uncharacterized IBP activity.

The genomic evidence for the presence of IBPs in intertidal species, paired with the behavioural and molecular limitations of intertidal invertebrates, led us to hypothesize that IBPs are important in achieving freeze tolerance in these species. Here, we sampled eight intertidal invertebrate species spanning three phyla in the intertidal zone in Vancouver, British Columbia, Canada. We measured all three classes of IBP activity, and tested for the occurrence of freeze tolerance in each species to see if IBP activity correlated with freeze tolerance. We predicted that freeze tolerant invertebrates in the intertidal would have IBP activity, whereas species that avoid freezing *via* behaviour would not. We also predicted that we would measure INP and IRI activity in each freeze tolerant species, and very little TH activity, since the latter is primarily used in physiological freeze avoidance.

## Methods

### Sample collection and study system

We sampled eight species of invertebrate during low tides at or below 1 m above chart datum (https://www.tide-forecast.com/locations/Vancouver-British-Columbia/tides/latest; details in Supplementary Methods). We then transported animals to the University of British Columbia Department of Zoology in zip-top bags or plastic containers, and immediately placed them in the −80 °C freezer if used for protein experiments. We tested the animals used in whole animal experiments within one hour of collection upon returning to UBC from the field site.

Additionally, we obtained individuals of *Acheta domesticus*, the common house cricket, from Jean-François Doherty (UBC Department of Zoology). These animals were prepared in the same manner as the intertidal organisms, and we used them throughout the assays as a non-freeze tolerant and non-intertidal invertebrate control. We used exclusively female crickets in all the experiments. Sample information in Table S1.

### Whole organism cold tolerance measures and supercooling points

All collections for this section were performed during the week of February 17, 2024 in the Vancouver, BC area (see supplemental methods for details). We subjected each animal to a cold event that approximated what they may face in the wild to measure cold tolerance. The cold exposure included a 10 minute hold at 11 °C, which was the air temperature of the warmest night of collecting. After the 10 minutes, the bath cooled at −1 °C per minute, until it reached −7 °C. We included the 18 minute ramp-down as part of the two hour exposure. At the two hour mark, we removed animals from cooling apparatus and put them in aquaria that matched the temperature and salinity of the collection site to simulate high tide. All following trials were given identical exposure conditions.

To measure the body temperature of each animal during the cold exposure, we placed a type T thermocouple (Pico Technology, UK) on the surface of the animal and held it in place with either sticky tac or Parafilm as necessary. The thermocouples were interfaced to a computer with a Picolog TC-08 which recorded temperature every 0.5 seconds. The temperature was controlled using a Haake AC200 controller on a Haake G50 cooling bath (Thermofisher; Burnaby, BC) filled with a ∼50:50 MeOH and water mixture. Depending on the size and mobility of each species, we placed individuals in either 1.5 mL microcentrifuge tubes, 15 mL plastic tubes, or sandwich bags within a larger zip-top bag. The tubes were placed in a custom aluminum bath head attached to the circulator, and the bags were placed directly in the circulating bath.

We determined the supercooling point (SCP) of each animal as the temperature immediately prior to the release of a large amount of heat due to the latent heat of crystallization (Lee, 2010, Figure S4). By the end of the cold exposure, we classified individuals as either frozen or unfrozen, as determined by the presence or absence of this exotherm. In the case of *H. oregonensis*, the 12 individuals exposed to −7 °C did not have their temperature tracked, and the SCP data came from a follow up experiment on March 6, 2024, where we were able to track body temperature.

After the cold exposure, we let the animals recover in an aquarium held in an incubator to control temperature. The water in each aquarium matched the temperature and salinity of each sampling site on the night of collection. Species collected at Stanley Park were put in 27 ppt salinity at 8°C, and species from Tower Beach were placed in 23 ppt at 7 °C (natural seawater from the UBC aquatic facility). The aquaria were oxygenated using air stones connected to an air pump. If individuals remained unfrozen by the end of the cold exposure, we separated frozen and unfrozen individuals in plastic containers with mesh siding. We monitored survival for each individual the week following the cold exposure. We defined survival as movement or response to physical stimulus. For *B. glandula*, we lightly poked the scutum with a fine probe and tested for resistance.

Control *A. domesticus* were also given a cold exposure under identical conditions to measure survival, but were recovered in a dry container at room temperature. Since none of those individuals froze during the −7 °C exposure, we exposed a second group of individuals to a continuous ramp down at −1 °C/min until all individuals froze to determine SCPs.

### Protein extraction

The protein extraction protocol we used was slightly modified from Box (2021). The homogenization buffer consisted of 50 mM ammonium bicarbonate (Thermo Fisher Scientific; Waltham, MA) with 0.1 mM phenylthiourea (PTU: TCI America; Portland, OR) and 0.1 mM phenylmethylsulfonyl fluoride (PMSF: EMD Millipore Corporation; Burlington, MA) at pH 8.00. We prepared a stock solution of 10 mM PTU and PMSF fluoride in 100% ethanol, and added to the samples immediately prior to use, alongside the 50 mM ammonium bicarbonate, to obtain a final concentration of 0.1 mM for PMSF and PTU. This buffer mix is hereafter referred to as the homogenization buffer.

We dissected as much tissue as possible from each animal and added it to 1.7 mL microcentrifuge tubes (Eppendorf Safe-lock; Eppendorf, Hamburg, Germany) with approximately 100 μL buffer per 100 mg of tissue. Samples were homogenized in a bead beater, then centrifuged at 16,100 × g for 10 minutes at 4 °C. Finally, we filtered the supernatant through a 0.45 μm (Millex HV [Merck Millipore; Cork, Ireland], Filtropur S [Sarstedt; Numbrecht, Germany]) syringe filter to eliminate any large nucleating contaminants.

We used the PierceTM BCA Protein Assay Kit following manufacturer’s instructions (Thermo Fisher Scientific; Waltham, MA) to determine the total protein content in technical triplicate for each sample. Each biological replicate for a species was standardized by dilution with additional homogenization buffer to the lowest protein concentration for that species (range of 2.9 mg/ml to 5.0 mg/ml; details in Table S2). We used samples at these standardized concentrations for droplet freezing and thermal hysteresis assays. All samples used in the IRI assay were diluted down to 0.25 mg/ml to match the concentration of BSA used in the assay. To isolate the effects of active protein on ice recrystallization inhibition and ice nucleation activity, we also separated each sample into two aliquots. One aliquot was sealed, and heat treated at 100-105 °C in an aluminum heat block for 10 minutes to denature proteins, while the other aliquot did not receive a heat treatment.

### Ice nucleation assay

We used a standard droplet freezing assay to determine ice nucleation activity for each of our samples. We slightly modified the routine cooling protocol (Ren et al., 2022; Whale et al., 2015) to suit our samples and more closely mimic an intertidal environment. Hydrophobic 18 mm circular glass cover slides (Hampton Research; Aliso Viejo, CA) were washed with Milli-Q water, blown with N_2_ until dry, and placed on a Grant Asymptote EF600 aluminum cooling stage (Grant Instruments; Shepreth, Cambridgeshire, England). We pipetted 1.0 μL droplets of protein extract onto the glass cover slips. For each freezing trial, we loaded 4 technical replicates for each of 5 biological replicates, plus 4 homogenization buffer droplets, for a total of 24 droplets per glass slide. This was repeated for the heat-treated aliquots for each species. The stage was covered with a Plexiglass cover, mounted with an LED strip to illuminate the stage and a video camera to record the droplets as they froze. Dry N_2_ was blown onto the stage at a rate of ∼0.2 L/m to prevent frost forming on the stage or the glass slides. The programmed cooling ramp was designed to decrease from room temperature (∼20 °C) to 0 °C over two minutes, held at 0 °C for two minutes, and then decreased by 1 °C/minute until all droplets were frozen. For analysis, the timestamped temperature log from the Grant Asymptote EF600 was cross-referenced with the time in the video where each droplet froze to determine nucleation temperatures.

### Ice recrystallization inhibition assay

We performed an ice recrystallization inhibition (IRI) assay following the protocol of Mitchell et al., 2015. The gold nanoparticles and samples were loaded in triplicate into a 96 well plate, along with positive (0.25 mg/mL BSA) and negative (50 mM ammonium bicarbonate) controls, also in triplicate, then read on a SpectraMax M2 Microplate Reader (Molecular Devices; San Jose, CA) on a spectrum from 400 nm to 700 nm in increments of 10 nm. The plate was then covered with a pipette tip box lid, then placed in a standard lab freezer set at −20 °C for 2 hours. We removed the plate from the freezer and allowed to thaw at room temperature until fully melted and was then read again at the same parameters. We calculated peak height in each spectrum by drawing a line between the values at 420 and 680 nm and taking the difference between that line and the measured absorbance value at 530 nm (see Supplemental methods, and Figure S1 & S2). We calculated IRI activity by subtracting the peak height before freezing from the peak height after freezing.

### Nanoliter osmometry

Thermal hysteresis measurements and ice shaping images were obtained using a nanoliter osmometer following manufacturer’s guidelines (Drori et al., 2013). We used a pipette puller to pull 0.4 mm I.D. glass capillaries (Drummond Scientific; Broomhall, PA) into micropipettes, and secured the capillary to a tube using adapters and sticky tack, fitted with a custom mouthpiece for injection. We used this device to suspend roughly 10 nL droplets of protein extract in oil (immersion oil B [Cargille laboratories, Cedar Grove, NJ]). A LabVIEW interface was used for temperature control (National Instruments), and droplets were visualized using a digital dissection microscope (Dino-Lite Edge 3.0 [Dino Canada; London, ON]). Once droplets were loaded, we lowered the temperature to −40 °C and waited until all droplets were frozen. After freezing, we raised the temperature back to determine the melting point of the solution. After increasing the temperature to a point where most of the ice had melted, we increased the temperature in increments of 0.005 °C to determine a precise melting point, defined as the highest temperature at which one remaining ice crystal barely shrinks. Once one crystal 30-45 nm in diameter was isolated at or near its melting point, we lowered the temperature by ∼0.01 °C (case-specific) so that it would neither grow nor shrink and started a 10-minute countdown. After the 10 minutes, the program was set to decrease temperature at 0.01 °C every 10 seconds. We recorded the last few seconds of the delay, in addition to the ramp-down to determine the crystal burst point / non-equilibrium freezing point of the sample, in addition to recording ice shaping activity if present. One droplet counted as one technical replicate. If there was no apparent ice shaping or hysteresis after testing three droplets (technical replicates) from two biological replicates, we classified the species as having no TH activity. In any individuals that showed ice shaping, we tested three technical replicates from all five individuals (biological replicates) of that species to examine whether results were consistent across biological replicates.

## Statistical analysis

All statistical analyses were completed in RStudio (R version 4.3.1; R Core Team, 2024). For the nucleation data, we took the average of the four technical replicates to get one nucleation temperature per biological replicate. For IRI data, we took the average of the three technical replicate wells for one measurement per individual. All calculations were performed with these mean values per biological replicate.

To calculate statistical significance among treatment groups (effect of heat treatment), we used a linear mixed effects model, with individual ID (biological replicate) as a random effect, and heat treatment as the fixed effect to predict nucleation temperature (nucleation data) or absorbance (IRI data). For the IRI data we ran an ANOVA on the linear mixed effects model for each species and adjusted the p-values using the “Hommel” method (Hommel, 1988).

We also calculated the differences in nucleation temperature or absorbance between active and denatured protein by taking the active protein mean value for one biological replicate and subtracting the denatured protein mean value of the same biological replicate. For the IRI data, we aimed to determine if active protein absorbance values fell above the activity of BSA. BSA acted as a “threshold” positive control, so any samples that that had an IRI activity at or above the BSA level were considered to exhibit IRI activity. To do this, we simply ran an ANOVA across all active protein samples and the positive BSA control.

## Results

### Do intertidal species have proteins with ice nucleation activity?

We detected protein-based ice nucleation activity in six out of eight intertidal species sampled, as measured by a significant difference in nucleation temperature between raw and heat-treated homogenate (Table 1). The highest nucleation temperature observed was in *My. trossulus*, followed by the only other bivalve sampled, *Ma. gigas* (Figure 1). We also investigated protein-based ice nucleation activity in seawater, finding that it was present, but that filtration (as we did with our biological samples) essentially removed this activity (Supplemental Results, Figure S3).

**Figure 1:**
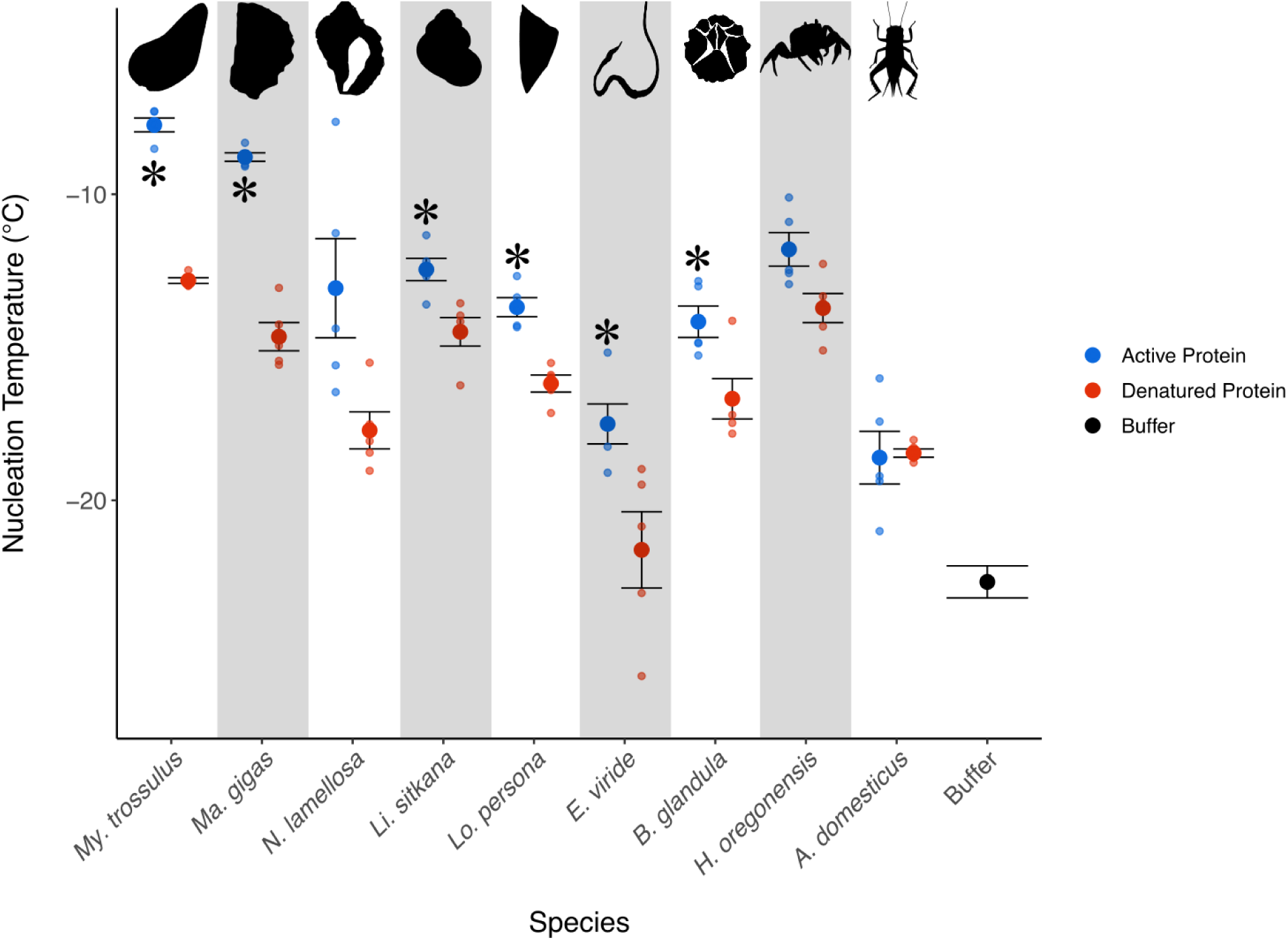
Nucleation temperatures of 1 µL droplets of homogenates of eight intertidal species, *A. domesticus*, and buffer. Large dots represent average nucleation temperature across biological replicates. Smaller dots represent values from biological replicates (mean of 4 technical replicates). Blue dots are samples filtered through a 0.45 μm filter, red dots are samples filtered and heat treated at 100 °C for 10 minutes to denature proteins. Bars represent standard error of the mean . * indicates significant difference (p<0.05) between active and denatured protein nucleation temperatures found via mixed-effects model after Hommel adjustment (statistics shown in table S3). Artists responsible for the silhouettes can be found in the references.

### Do intertidal species have proteins that produce ice recrystallization inhibition (IRI)?

We assessed IRI activity using a gold nanoparticle-based assay. We found that all active protein absorbance values for all species were above the value from BSA, the positive control, except for *H. oregonensis* (Figure 2). However, only *My. trossulus*, *Ma. gigas*, and *L. persona* had significantly lower IRI activity when proteins were denatured, as assessed by an ANOVA and following Hommel adjustment (Table 2). Strangely, there was a significant rise in IRI activity measured from *H. oregonensis* homogenate when proteins were denatured. For all other species, including the non-intertidal control *A. domesticus*, there was no significant difference in IRI activity between active and denatured protein, but all values remained at or above the level of BSA indicating other non-protein substances likely causing IRI.

**Figure 2:**
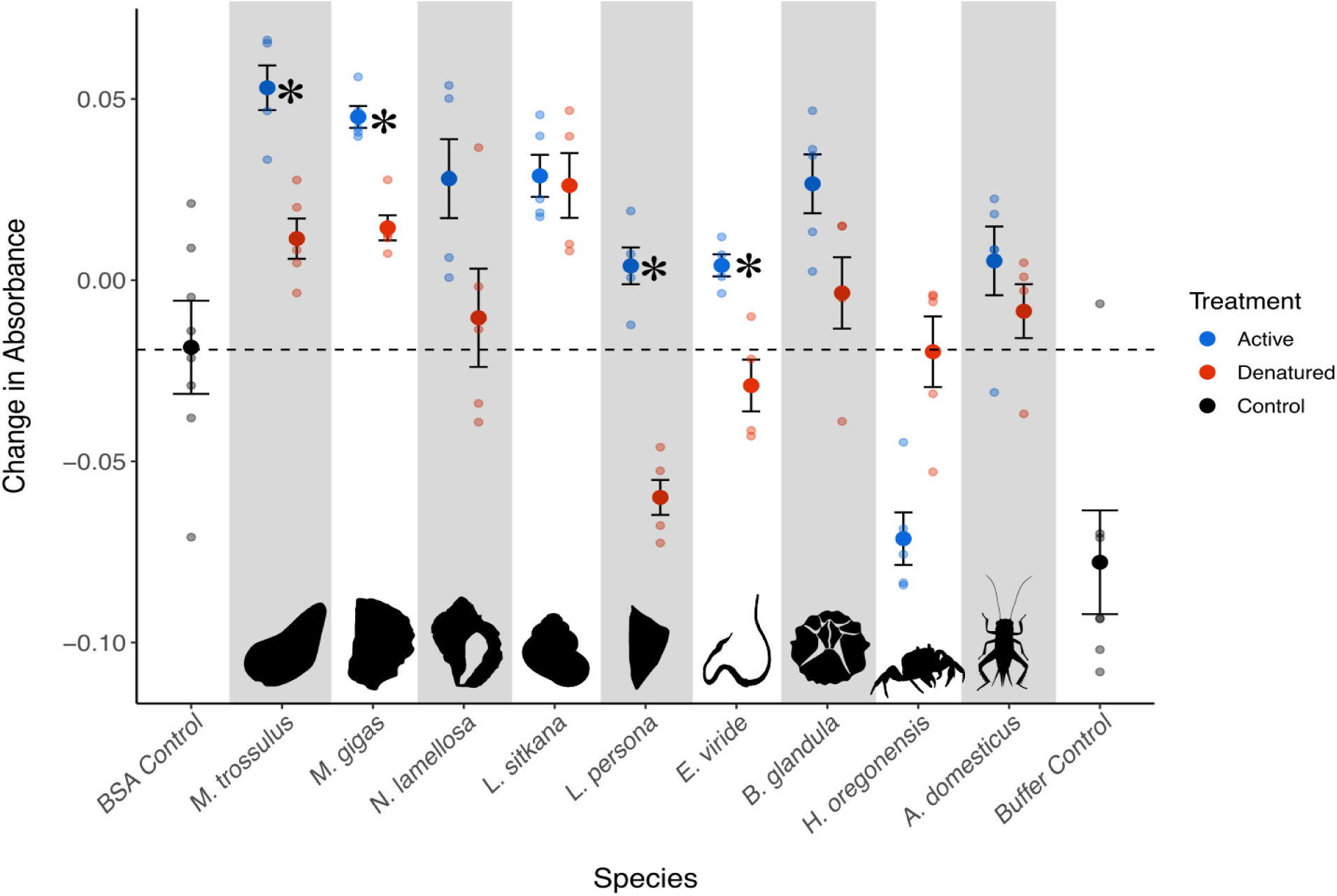
Change in absorbance values after freeze-thaw cycle for active and denatured protein samples in intertidal species, *A. domesticus*, and controls, with higher change in absorbance indicating higher IRI activity. All samples were standardized to 0.25 mg/ml total protein. Three wells of positive (BSA) and negative (Buffer) controls were run on each plate alongside samples, and the average values (small black dots) were pooled across trials to calculate overall mean values (large black dots). Blue dots are unboiled samples, red are boiled (denatured). Small dots represent mean of technical triplicates for each biological replicate, large dots are means of biological replicates (n = 5). Dotted black line is at y= − 0.0192, the mean of all BSA trials. * indicates significant decrease (p<0.05) in absorbance values if proteins are denatured, in individual mixed-effects models following Hommel adjustment (Table S4).

### Do intertidal species have proteins that produce thermal hysteresis?

Thermal hysteresis (TH) is typically defined as the difference between the melting point and the non-equilibrium freezing point – the point at which a burst of crystal growth occurs (Drori and Stevens, 2024). Using this definition, none of the species tested displayed any TH, since they all lacked a crystal burst point. However, we did find a few species that displayed ice shaping, which is generally a diagnostic feature of TH proteins, since ice growth in the absence of TH proteins will occur in circular sheets (DeLuca et al., 1996). We observed a variety of ice crystal shapes in our samples, some of which have been observed in the literature, while some appear novel (Figure 3). Species that showed ice shaping in all five biological replicates included *My. trossulus*, *Ma. gigas*, and *H. oregonensis*. All of these taxa displayed a mix of ice shaping behaviours, with the most common shape occurring being hexagonal. These hexagonal ice planes have been observed many times in the literature, and are often indicative of concentrations of TH proteins too low to cause significant TH (Lorv et al., 2014; Rahman et al., 2019). The “hourglass” shape observed in *My. trossulus* and *H. oregonensis* is reminiscent of the shape observed during the explosive burst of growth seen in hyperactive AFPs (Bar-Dolev et al., 2012), the only difference being our samples displayed no burst, and instead showed slow and steady growth towards this shape.

**Figure 3:**
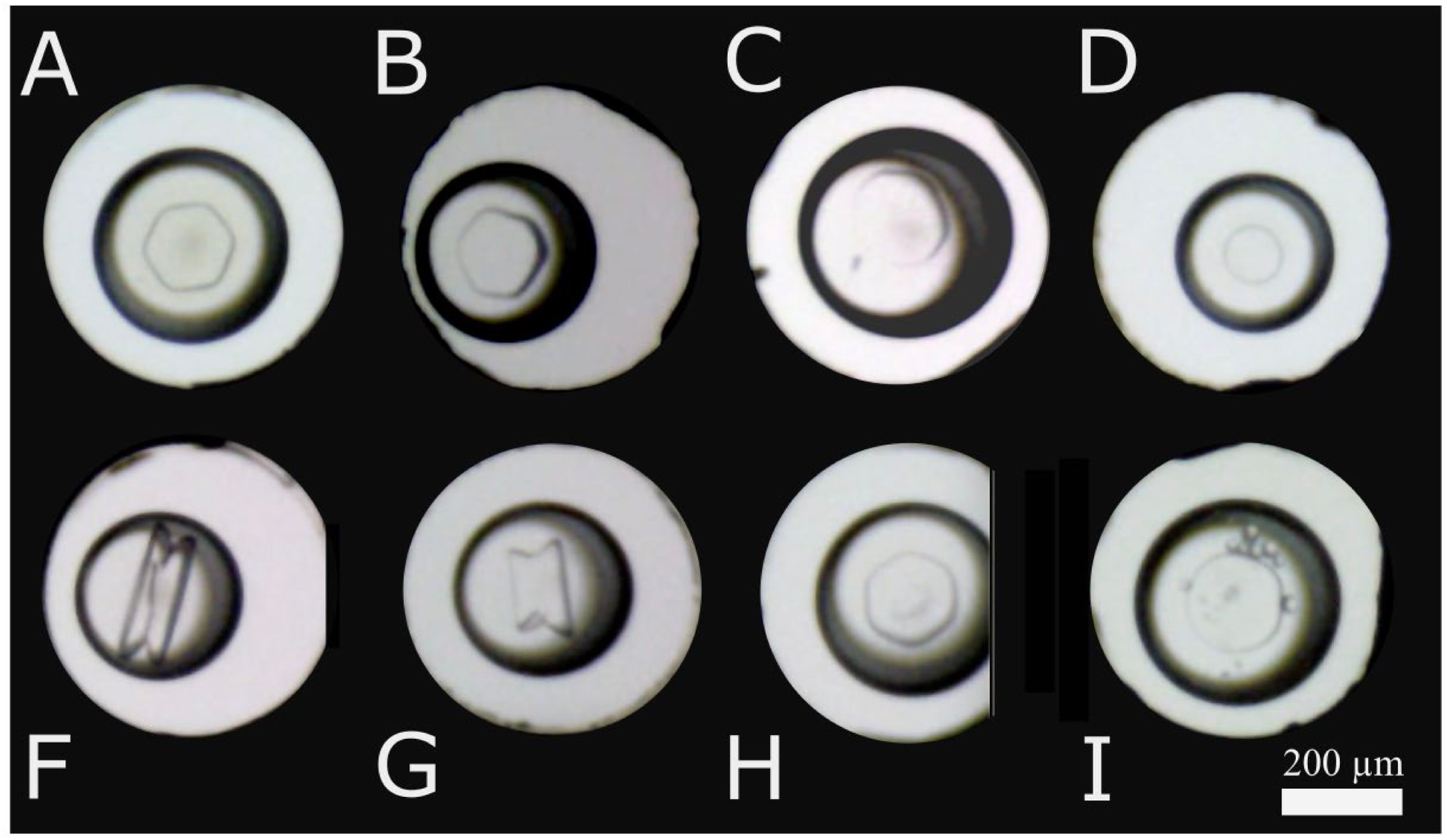
Examples of ice shaping in intertidal species. Ice shaping shown in *H. oregonensis* (A & F), *My. trossulus* (B & G), *Ma. gigas* (C), *Lo. persona* (H). Buffer (I) and *E. viride* (D) show no ice shaping. Protein extracts from intertidal species in a nanoliter osmometer, captured with a DinoLite® digital microscope. Samples were frozen at −40 °C, then brought up to their melting point to isolate one ice crystal ∼30-45 µm in diameter. Single crystals were held ∼0.01 °C below their melting point for 10 minutes, then the temperature decreased by 0.01 °C every 10 seconds.

### Does IBP activity impact survival or behaviour on a whole animal scale?

Data for the following sections were obtained from whole animal cold exposures to −7 °C for 2 hours. In the case of *N. lamellosa*, some individuals did not experience −7 °C due to flaws in the experimental set up. Two individuals only reached ∼-3.5 °C, and one only −5 °C; none of these individuals froze. We found that most intertidal species freeze at relatively high sub-zero temperatures, with all species freezing between −2.61 and −5.86 °C. The highest SCPs were in *Ma. gigas* (−2.61 ± 0.293 °C) and lowest were in *L. sitkana* (−5.86 ± 0.121 °C) and *H. oregonensis* (−5.86 ± 0.209 °C; Figure 4).

**Figure 4:**
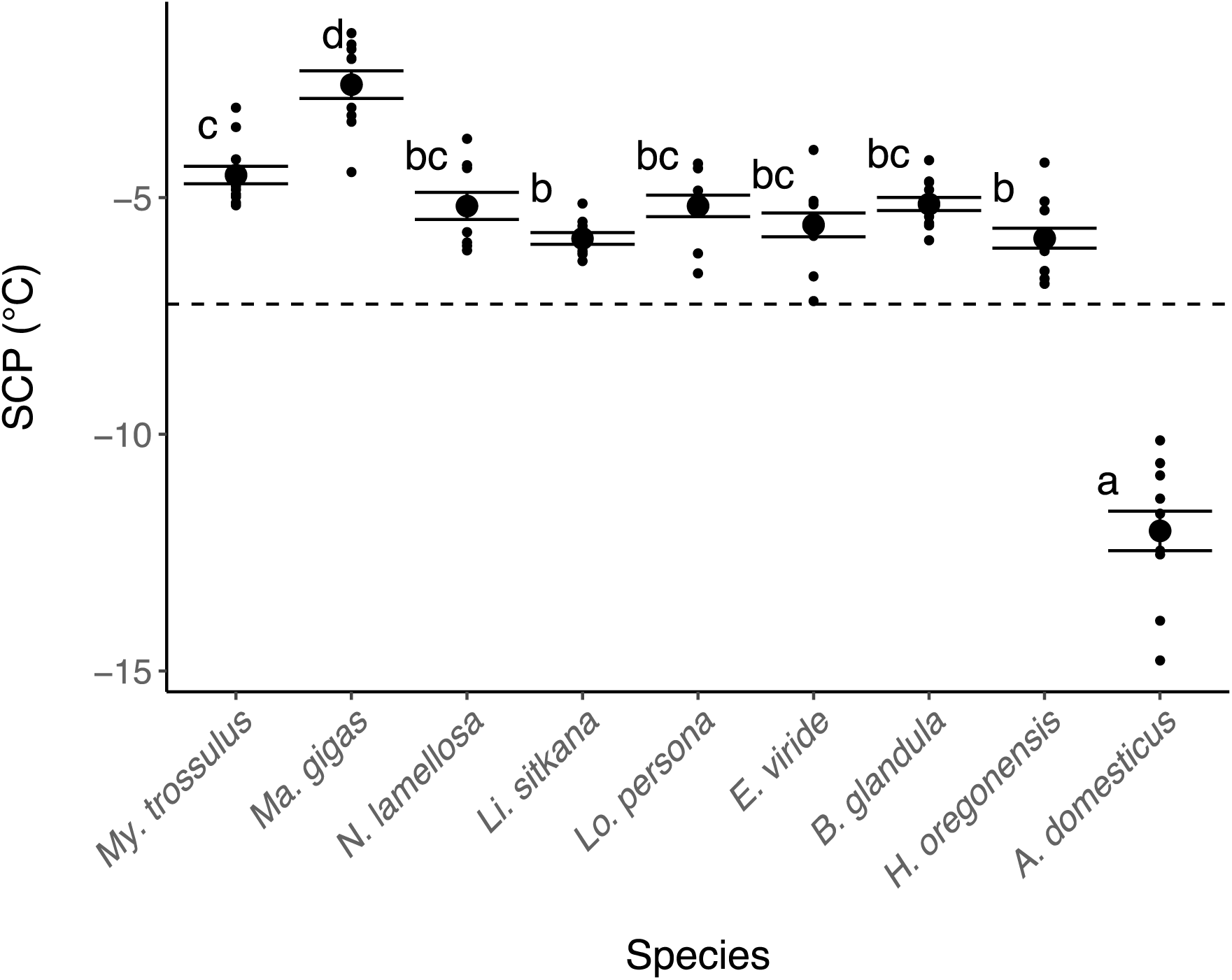
Whole organism supercooling points for each intertidal species. Data was collected from the cold exposure trials. 12 individuals were tested for each species. SCPs were included for each frozen individual unless the SCP was not clear, the thermocouple detached during the trial, or the SCP occurred after the temperature ramp. In order of species displayed, N = (12, 10, 9, 10, 10, 12, 12, 11, 11). Small dots represent individual animals, large dots represent means of biological replicates. Error bars are standard error of the mean. Matching letters represent no statistical difference (p<0.05) in a Tukey posthoc test using emmeans package following ANOVA.

### Survival after cold exposure

For most of the species studied, there was a clear exotherm during the cold exposure, meaning the individual experienced freezing (Figure S4). We classified species that recovered after a freezing event as freeze tolerant (Table 3). For *Ma. gigas*, there were a few individuals in which we could not identify a clear exotherm. Instead of a regular SCP, once the individual hit a certain temperature, it remained stable for 20 minutes to 1 hour, before continuing to decrease (supplemental Figure S4). Only one *N. lamellosa* that froze survived, leading us to classify the species as partially freeze tolerant. No *H. oregonensis* survived the two-hour cold exposure.

**Table 3:** Freezing and survival data for each species after a two hour exposure at −7 °C. Temperature was lowered by −1 °C/min from 11 °C, and body temperature was tracked using a type T thermocouple, connected to a computer running PicoLog software recording every 0.5 seconds, to determine if individuals froze.

| Species | Status | Number<br>alive<br>following<br>cold<br>exposure | Number<br>dead<br>following<br>cold<br>exposure | Likely cold<br>tolerance<br>strategy |
| --- | --- | --- | --- | --- |
| *My. trossulus* | Frozen | 12 | 0 | Freeze tolerance |
|  | Unfrozen | 0 | 0 |  |
| *Ma. gigas* | Frozen | 12 | 0 | Freeze tolerance |
|  | Unfrozen | 0 | 0 |  |
| *N. lamellosa* | Frozen | 1 | 8 | Partial freeze<br>tolerance |
|  | Unfrozen | 3** | 0 |  |
| *Li. sitkana* | Frozen | 11 | 0 | Freeze tolerance |
|  | Unfrozen | 1 | 0 |  |
| *Lo. persona* | Frozen | 10 | 0 | Freeze tolerance |
|  | Unfrozen | 2 | 0 |  |
| *E. viride* | Frozen | 5 | 5 | Partial freeze<br>tolerance |
|  | Unfrozen | 1 | 1 |  |
| *B. glandula* | Frozen | 10 | 2 | Freeze tolerance |
|  | Unfrozen | 0 | 0 |  |
| *H. oregonensis* | Frozen | 0 | 12* | Partial freeze-<br>tolerance |
|  | Unfrozen | 0 | 0 |  |
| *A. domesticus* | Frozen | 0 | 0 | Chill susceptible<br>or freeze-<br>avoidance |
|  | Unfrozen | 12 | 0 |  |

**\*We did not track body temperature for the *H. oregonensis* cold exposure, but each individual was visibly frozen and brittle. We followed up by tracking body temperature for SCP measurements where 50% of individuals recovered after being frozen for ~10 minutes at -7 °C.**
**\*\*Unfrozen individuals were only exposed to -3.5 °C (N= 2), and -5 °C.**

However, when we measured SCPs in a follow-up experiment, we removed each animal from the cooling bath 5-10 minutes after freezing, and left them for recovery in an aquarium. Half of these animals survived, leaving us to classify them as partially freeze tolerant as well.

## Discussion

Here we hypothesized that ice binding proteins, specifically those causing IN and IRI activity, play an important role in achieving freeze tolerance in intertidal invertebrates. We also predicted we would find very little TH activity, since freeze avoidance is not a common strategy in intertidal invertebrates. We found evidence that IBPs, particularly those causing IN activity, are widespread in the temperate Pacific intertidal. No strong TH was detected in any species, but we did find evidence that TH proteins are present, just at very low concentrations relative to the whole organism level. We also found evidence that there is IRI activity attributable to proteins in a few intertidal species. We largely found that species that displayed the highest survival after freezing showed evidence of INPs, indicating INPs may be important for achieving freeze tolerance in intertidal invertebrates (Table S5). This first broad survey of IBPs across a habitat indicates that species in the temperate intertidal zone contain many novel ice binding proteins, and adds significantly to our knowledge of stress tolerance in this habitat.

### Ice Nucleation

Our results suggest that INPs, as evidenced by the ablation of nucleation activity when proteins were denatured in six out of eight intertidal species assayed, may be widespread in temperate intertidal species. Of these species, IBP activity has only ever been investigated in *My. trossulus*, using chemical treatments to denature protein (Box, 2021). Instead of chemical treatments, we decided to simply heat-treat all samples to denature proteins. Heat treatment has been shown to significantly reduce the nucleation temperature of INPs, but not completely eliminate it (Hansen et al., 2023). Ice nucleation is believed to be an important aspect of freeze tolerance, however relatively few biological ice nucleators are characterized (Gill et al., 2024; Storey and Storey, 1988; Zachariassen and Kristiansen, 2000). Here we add significantly to the known animal species that contain biological ice nucleators.

### Ice shaping, TH, and IRI

The overwhelming majority of IBP literature has historically grouped IRI proteins and TH proteins under the term “antifreeze protein” (AFP). This is largely because these two activities are strongly connected, with all known TH proteins also having IRI activity (Olijve et al., 2016; Yu et al., 2010). In other words, all AFPs have both TH and IRI in different contexts: TH before ice formation and IRI after ice formation. *In vivo*, organisms likely utilize AFPs in different ways based on their cold tolerance strategy. Freeze-tolerant species likely use AFPs for IRI and IN, and freeze-avoidant organisms utilize TH activity (Bar Dolev et al., 2016).

We found no measurable TH in any species. However, there was evidence that TH proteins may have been present at low concentrations in some of the species. We found that *My. trossulus*, *Ma. gigas,* and *H. oregonensis* homogenates all showed ice shaping in most or all replicates tested from each individual (Figure 3). We also found evidence of ice shaping in *Lo. persona*, but to a much lesser extent, only appearing in about half of the technical replicates tested. There are a few documented cases of ice-shaping activity without measurable TH in cases of low protein concentration (DeLuca et al., 1996; Lorv et al., 2014; Rahman et al., 2019). For example, an IBP from snow-mold fungus can achieve its maximum IRI activity at 1 μM, but needs concentrations of over 500 μM for any measurable TH. In this initial survey, we chose to homogenize whole body tissues from all species as we did not have a specific hypothesis for which tissues may express IBPs in each species. However, if TH proteins were concentrated in certain intracellular compartments or specific organs, this method of preparation may have diluted TH proteins to a concentration too low to measure activity. At low concentrations of IBPs where IRI can be measured, the ice is often shaped into hexagonal planes (Rahman et al., 2019), like those observed in our experiments.

We also observed non-hexagonal ice crystal growth, which could be evidence of entirely novel motifs of IBPs, or just evidence of incomplete ice binding (Figure 3). We found *My. trossulus Ma. gigas*, and *Lo. persona* displayed characteristic ice shaping of IBPs, in addition to having protein-based IRI activity, while *H. oregonensis* showed ice shaping without IRI activity. There is a slight caveat in that the IRI results from *H. oregonensis* are unusual, with the denatured protein showing more IRI than the active protein samples (Figure 2). We believe the cause of this abnormality is due to protein interacting with the gold nanoparticles, causing aggregation that did not relate to ice formation. Therefore, we do not believe we can adequately assess IRI capacity in *H. oregonensis* with this method. While the ice shaping results show clearly that this species expresses an IBP, whether or not that IBP causes IRI cannot be assessed with our data.

There are two documented examples of TH activity in intertidal invertebrates, however both cases found activity in extracted haemolymph, instead of whole-animal homogenate. The blue mussel and congener of *My. trossulus, My. edulis,* has extremely low TH, and the Antarctic nemertean *Antarctonemertes validum,* has a moderate TH of 1.4 °C (Theede et al., 1976; Waller et al., 2006). There is also some evidence that the Manila clam, *R. phillippinarum,* expresses an IBP similar to type II AFPs from fish, although the TH potential of this IBP has not been investigated (Dong et al., 2022). There is additional genomic evidence that many intertidal invertebrates have IBP-like sequences that show high similarity to TH proteins in fish, but the activity of these proteins has not been measured (Box et al., 2022). We believe it is likely that TH proteins are present, but at very low concentrations in each of the species with ice shaping. By homogenizing the whole organism, we may have diluted any TH proteins to a level where the activity cannot be measured, especially if the TH proteins were localized in the haemolymph.

Alternatively, TH proteins may be expressed only in particular organs, as was the case with *R. phillipinarum*, or in intracellular spaces, and extracting haemolymph would also not adequately capture TH.

### Whole organism implications

We predicted that not only that IBPs would be widespread, but that IBP type and presence would relate to cold tolerance strategy. We found that, of the species that are highly freeze-tolerant, most have evidence for INPs. Those species were *My. trossulus, Ma. gigas, Lo. persona, Li. sitkana,* and *B. glandula*. We also found evidence for INPs in *E. viride,* but this species had lower survival following cold exposure. Of these species, *My. trossulus,* and *Ma. gigas* also showed ice shaping, indicating non-nucleating IBPs in addition to INPs. IBPs also appear to be present in species that are relatively poor at surviving the cold, like *H. oregonensis*. The IRI data for this species was highly unusual (Figure 2), making conclusions difficult, but the presence of ice shaping activity indicates that this species does produce an IBP.

The ability to behaviourally thermoregulate does not appear to play a major role in IBP presence. Both *H. oregonensis* and *E. viride* are highly mobile and can retreat into warmer microhabitats if needed. *Hemigrapsus oregonensis* is almost exclusively found underneath rocks at low tide, and nemerteans have been shown to preferentially emerge at climatically-favorable low-tides (Caplins et al., 2012). We found that *E. viride* contains INP activity, while *H. oregonensis* contains strong ice shaping activity. Although there was no measurable TH in *H. oregonensis*, and it showed no additional signs of IBPs, the ice shaping activity is evidence of IBP presence in this species. To further study the effect of behavioural thermoregulation on IBP evolution would require a targeted study comparing closely-related taxa.

We found intertidal position was generally a good predictor of freeze tolerance, with the highest shore intertidal species having the highest survival, but we did not find a pattern between IBP type and shore position. *Mytilus trossulus*, *Ma. gigas,* and *Lo. persona* can all be found very high in the intertidal, and were all found to have protein-based IRI activity. By contrast, *B. glandula* and *Li. sitkana* can also be found very high in the intertidal and were not found to have IRI activity. All these high intertidal species were found to have INP activity. In the low intertidal, *N. lamellosa* is left emersed for relatively short periods of time. We found high mortality in this species after the cold exposure and also INP activity, which presents a puzzle. It is possible that they do not express endogenous INPs, and that any nucleating activity stems from their environment or diet.

In conclusion, we found that IBP activity is widespread in the temperate Pacific intertidal, with all intertidal species sampled showing signs of at least one IBP activity. The source of these IBPs is still unknown, but there is genomic evidence that *My. trossulus* and *Ma. gigas* express endogenous IBP genes. Before this study, there were only six intertidal species with documented IBP activity. Here we more than double the number of species with evidence for INP activity, in addition to adding species that contain IRI and ice shaping activities. This is clear evidence that the intertidal is indeed an ecosystem that fosters the evolution of IBPs, and adds significantly to our knowledge about how these proteins evolve.

## Acknowledgements

Data and code is available at (RELEASED ON PUBLICATION).

## Acknowledgements

NM was supported by a UBC Faculty of Science grant, while KEM and AKB are supported by individual NSERC Discovery grants. Thank you to Soleil Worthy and Di Chen in the Bertram Lab for the training and assistance with the droplet freezing procedure, and to Isaiah Box for the training and assistance with the nanoliter osmometer.

