## Supplemental data for "ICE BINDING PROTEIN ACTIVITY IS COMMON AMONG TEMPERATE PACIFIC INTERTIDAL INVERTEBRATES"

**Supplemental methods**

We sampled all animals at one of two locations in Vancouver, British Columbia, Canada. Our two collection sites provided access to different species due to their distinct ecological contexts. Tower Beach (49.273681 °N 123.257514 °W) is located right near the mouth of the Fraser River, so the salinity of the water fluctuates wildly between seasons due to changes in river flow. In the summer months, the salinity can drop to below 10 ppt, and sits between 25-30 ppt for most of the winter (Covernton and Harley, 2020). This habitat is only suitable for animals that can withstand these salinity changes. All *My. trossulus, Ma. gigas, B. glandula, H. oregonensis, Li. sitkana,* and *E. viride* collections were from Tower Beach. The other collection site was in Stanley Park near the “Girl in Wetsuit” statue (49.302668 °N 123.126293 °W). This site maintains a more stable salinity (~25 ppt) throughout the year, and can occasionally drop below 20 ppt in the summer (Covernton and Harley, 2020). This site allowed us to sample additional species that we could not find at the Tower Beach location. We collected *N. lamellose* and *Lo. persona* at this site. We pried limpets off of rocks with a credit card, and all other species were collected by hand (Table S1). We did not record sex for any intertidal species sampled, except for *H. oregonensis*, which displays clear sexual dimorphism. We used one female and four male crabs for the IBP assay work, and an equal mix of males and females for the whole organism experiments. All sampling occurred in accordance with Department of Fisheries and Oceans scientific collection permits XMCFR 35 2022 AMD02 and XMCFR 49 2023. At the sampling site, we took a measure of the salinity, sea water temperature, and air temperature with a YSI Pro30 Conductivity meter (Xylem Inc; Yellow Spring, OH, USA).

We used different dissection techniques for each species to maximize the amount of tissue collected from each animal. For example, we used a scalpel to scrape as much tissue from the mollusc shells as possible, whereas for *H. oregonensis* we used a scalpel to cut open the shell and used fine probes to scrape out tissue. In cases where we needed to break open shells, for example with *Li. sitkana*, we took care to avoid adding shell particles to the microcentrifuge tubes.

To ensure that no more than 600 mg of tissue was added to each tube, larger animals were split up between multiple tubes. A scoop of 0.9-2.0 mm stainless steel beads (Next Advance; Troy, NY), roughly equaling the mass of tissue, was added with the tissue. Buffer quantity varied by species, with approximately 100 μL buffer per 100 mg of tissue. We never used less than 100 μL of buffer, even in cases of extremely low tissue mass. We homogenized tissue in a Storm 24 Bullet Blender (Next Advance; Troy, NY) at maximum speed for 4 minutes. Tissue was re-homogenized if there were still large pieces visible after the initial 4 minute period. We then centrifuged the samples at 16,100 × g for 10 minutes at 4 °C. Supernatant was removed and put in a new tube. If individual animals were initially divided between multiple tubes, we pooled the supernatant into a 15 or 50 mL centrifuge tube. The pellet was then resuspended in homogenization buffer and put back into the Bullet Blender for another 2-4 minutes. Centrifugation was repeated, and the supernatant pooled with the homogenate from the previous round. We ran another round of bullet blending and centrifugation if the supernatant from the second round was still very dark or cloudy. The resulting pooled sample contained total soluble protein extracted from the entire animal.

Table S1 Taxonomic information for each species collected. Each species and genus name appears as accepted on the World Register of Marine Species (<https://www.marinespecies.org/index.php>). Collection dates include animals collected for protein analysis^1^, and whole animal analysis^2^.

**H. oregonensis* SCPs were taken on 03/06/2024 since temperature was not tracked during the 2-hour cold exposure.

| **Name** | **Phylum** | **Class** | **Location Collected** | **Dates**  **Collected**  **(DD/MM/YYYY)–** | **Shore height where animals were collected** |
| --- | --- | --- | --- | --- | --- |
| *Mytilus trossulus* | Mollusca | Bivalvia | Tower Beach | ^1^ 08/12/2022  ^2^ 17/02/2024 | Middle-High |
| *Magallana gigas* | Mollusca | Bivalvia | Tower Beach | ^1^ 08/12/2022  ^2^ 20/02/2024 | Middle |
| *Littorina sitkana* | Mollusca | Gastropoda | Tower Beach | ^1^ 15/03/2023  ^2^18/02/2024 | High |
| *Nucella lamellosa* | Mollusca | Gastropoda | Stanley Park | ^1^ 06/01/2023  ^2^19/02/2024 | Low |
| *Lottia persona* | Mollusca | Gastropoda | Stanley Park | ^1^ 16/03/2023  ^2^19/02/2024 | High |
| *Emplectonema viride* | Nemertea | Hoplonemertea | Tower Beach | ^1^ 17/02/2024  ^2^17/02/2024 | Middle |
| *Balanus glandula* | Arthropoda | Thecostraca | Tower Beach | ^1^ 08/12/2022  ^2^ 17/02/2024 | Middle-High |
| *Hemigrapsus oregonensis* | Arthropoda | Malacostraca | Tower Beach | ^1^ 15/03/2023  ^2^ 20/02/2024* | Low-Middle |

#### DNA Barcoding

After investigating the small *Littorina spp*. snails we collected, we discovered that we had collected three different species, with *Li. plena* and *Li. scutulata* having near-identical shell morphologies. To distinguish these two species from each other, we used extracted DNA from one tentacle of frozen specimens using Qiagen DNEasy Blood and Tissue kit following manufacture’s protocol with minor modifications (Qiagen; Hilden, Germany). We used a Kapa Taq Ready Mix PCR kit according to package directions (KapaBiosystems; Cape Town, South Africa). Attempts to distinguish these species remained unsuccessful, so we only included *Li. sitkana,* which can easily be distinguished from its two local congeners using shell morphology. We also extracted DNA from the *Mopalia spp.* and *Emplectonema viride* specimens used for protein analyses to identify both at a species level. Samples were amplified using Folmer primers (Folmer et al., 1994), and sent in for Sanger sequencing at the UBC Sequencing and Bioinformatics Consortium. We checked the resulting FASTA sequences against the NCBI database, and viewed the trace data in Benchling (www.benchling.com) to make sure base pair calls were clear.

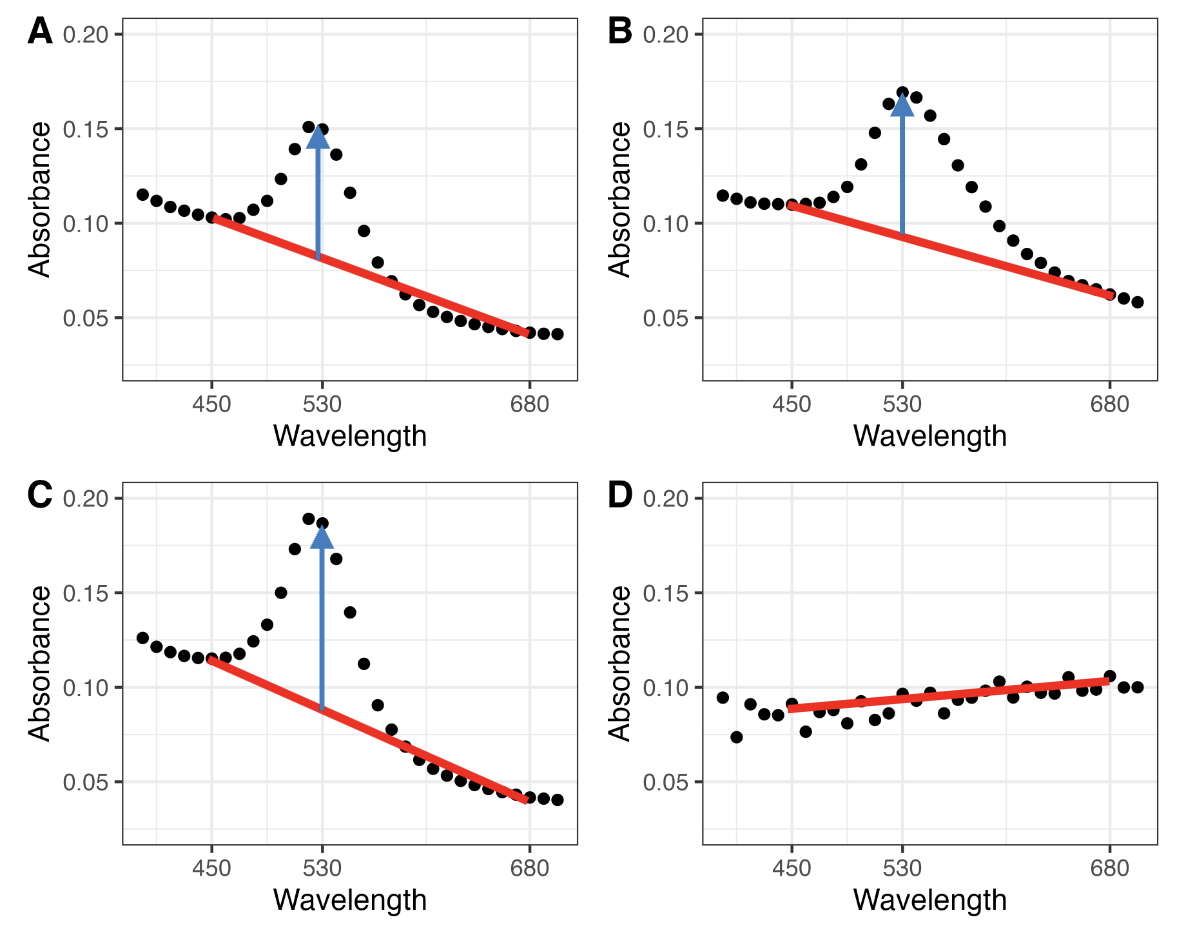

Figure S1. Graphical representation of IRI data analysis. Example spectra of threshold IRI activity in positive control BSA (0.25 mg/ml) before (A) and after (B) freeze-thaw cycle. Negative control 50 mM ammonium bicarbonate before (C) and after (D) freeze thaw cycle. Red line is drawn between values at 420 and 680 nm, blue arrow shows peak height at 530 nm. IRI activity is measured by taking the difference in peak height before and after freeze-thaw cycle.

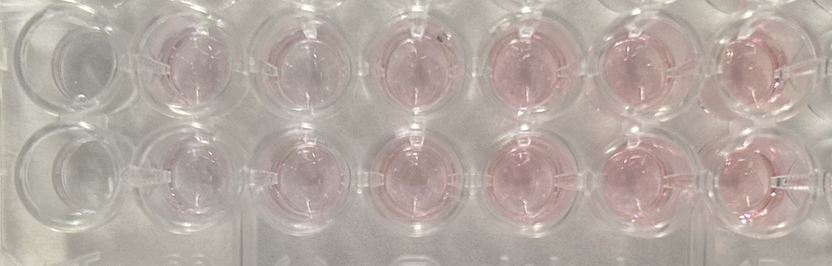

Figure S2: Color change after freeze-thaw cycle in a standard curve of bovine serum albumin (0, 0.25, 0.5, 1, 2.5, 5, 10 mg/ml, left to right).

#### Seawater freezing

We measured the nucleation potential of untreated and 0.45 μm filtered seawater collected at Tower Beach in two different assays to compare the results to both whole animal and droplet freezing assays. Some of the species, particularly the bivalves, retained a significant amount of seawater during the SCP trials, and in the original tissue dissections. For this reason, we wanted to investigate the nucleating potential of that seawater to differentiate the effects of any animal IBPs vs any contaminants from the environment. We investigated the nucleation potential using the droplet freezing procedure as described above, on untreated, filtered, and filtered and boiled samples, which matched the treatments that the animal homogenate received. Additionally, we ran an assay that mimicked the whole animal SCP assay using 1.0 mL samples of seawater. We filled three 1.5 mL microcentrifuge tubes with filtered sea water, and three tubes with unfiltered seawater, placed them in the same cooling blocks used in the SCP trial, and lowered the temperature at 1 °C/min. We attached thermocouples to the outside of the tubes to record temperature. We ran this test three times in a row, after each tube had fully melted, giving nine SCPs for both filtered and unfiltered seawater tubes.

Table S2: Total protein concentration for each species homogenate used in ice nucleation and thermal hysteresis assays. Species include all nine intertidal species sampled, and cricket control.

| **Species** | **Protein concentration (mg/mL)** |
| --- | --- |
| *My. trossulus* | 3.83 |
| *Ma. gigas* | 3.92 |
| *N. lamellosa* | 4.52 |
| *Li. sitkana* | 2.94 |
| *Lo. persona* | 5.00 |
| *E. viride* | 3.78 |
| *B. glandula* | 4.28 |
| *H. oregonensis* | 5.00 |
| *A. domesticus* | 5.00 |

**Supplemental Results**

We investigated how seawater from one of the sample sites would behave in some of the assays conducted, compared to the results of how animal tissue that contains the same seawater. We collected seawater from the Tower Beach site to use in a “whole animal” SCP assay, as well as the droplet freezing assay to test for nucleators. In the SCP-style assay, we filled microcentrifuge tubes with 1 ml of 0.45 μm filtered or unfiltered seawater. This volume roughly approximated the amount of water held in the shells of an average *My. trossulus* individual collected*.* efound that the unfiltered seawater froze at -7.25 ± 0.23 °C, and the filtered seawater froze at a significantly lower temperature at -10.40 ± 0.49 °C (p < 0.001, Figure 2.11A). For comparison, the lowest SCP in an intertidal species was from *H. oregonensis* at -5.86 ± 0.21 °C. Likewise, in the droplet freezing assay we found that filtration significantly decreases the nucleation temperature. The unfiltered seawater nucleated at -12.90 ± 0.54 °C, whereas filtered seawater nucleated much lower, at -23.20 °C ± 0.61 °C (p < 0.001**)**, indistinguishable from buffer droplets.

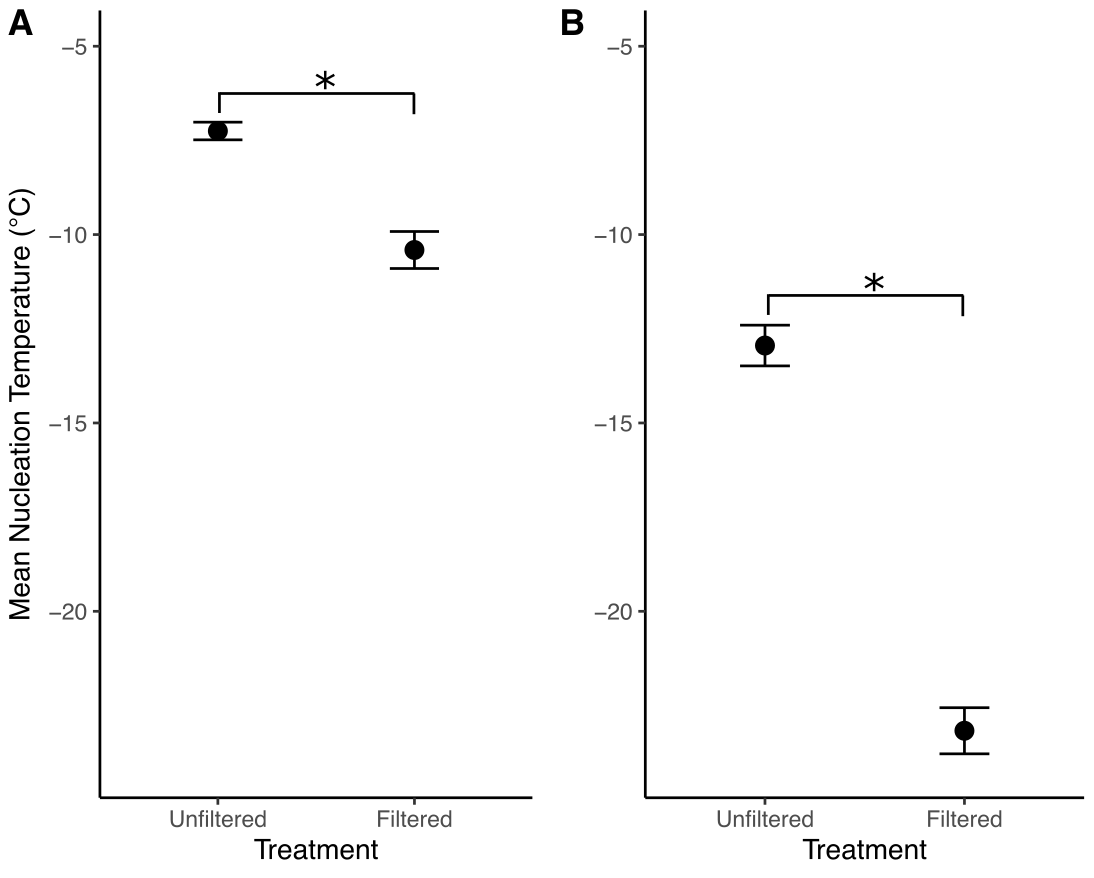

**Figure S3: Nucleation temperatures for 1.0 mL (A) and 1.0 μL (B) of filtered and unfiltered seawater. A) Nucleation temperature of 1.0 ml of seawater from Tower Beach, unfiltered and filtered through a 0.45 μm syringe filter. B) Nucleation temperature of the same seawater but tested in 1.0 μL droplets in the droplet freezing assay described in methods. Both assays had a cooling rate of -1 °C/min. * indicates significance found via t-test (p < 0.001 in both cases)**

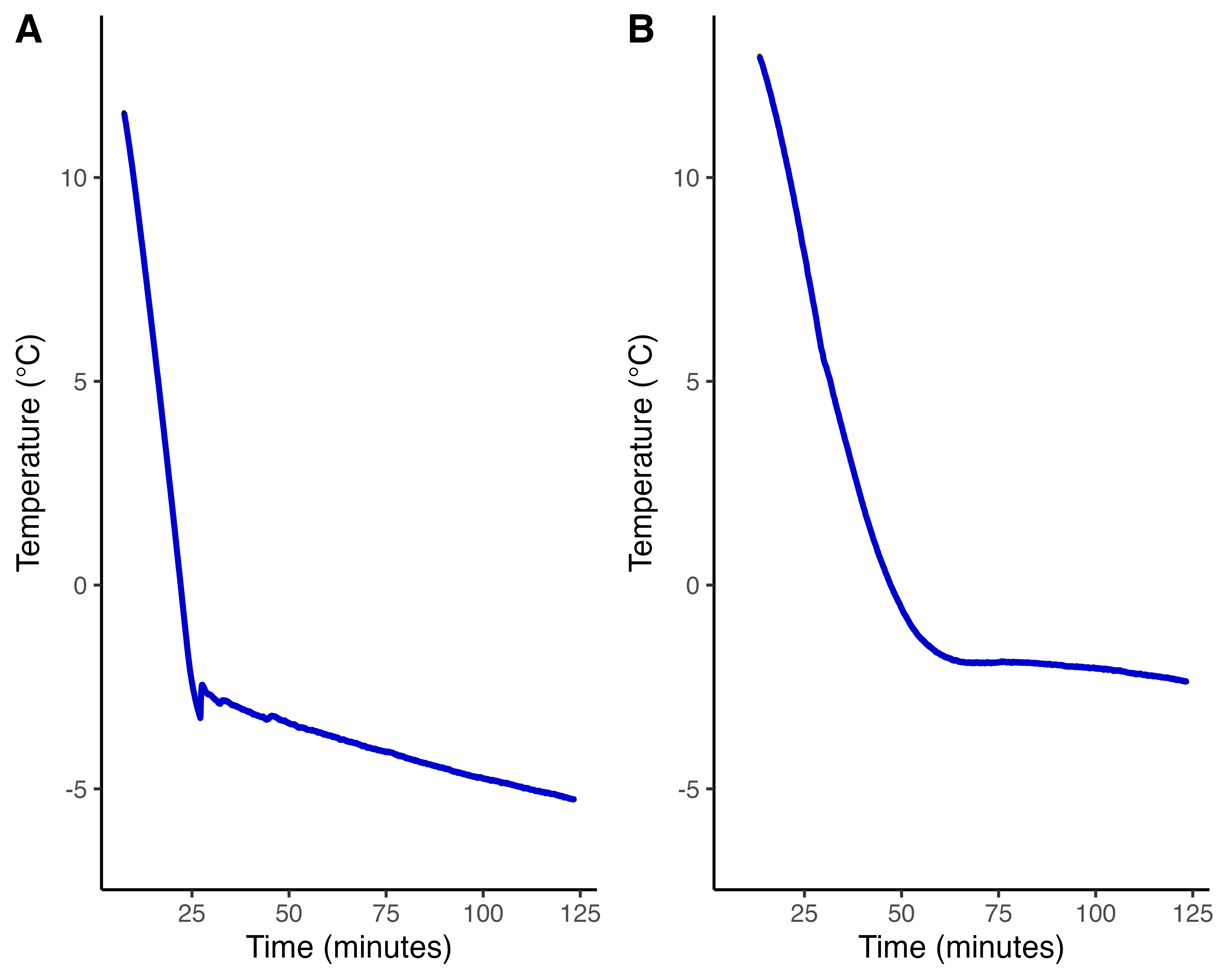

Figure S4: Temperature traces of two *M. gigas* individuals during cold exposure.

(A) Example of a “normal” SCP. (B) Example of a flattening of the temperature, showing a slow and prolonged freeze. No SCP value recorded for individuals like this. Ambient temperature started at 11 °C, and was lowered to -7 °C at 1 °C/min.

Table S3: Effect of heat denaturation treatment on nucleation temperature of 1 µL droplets of homogenate in intertidal species. F statistics and P-values are from an ANOVA on each individual mixed-effects model for each species, with individual ID as a random effect. Bolded values indicate statistical significance after a Hommel adjustment (p<0.05).

| **Species** | **Effect of boiling on nucleation temperature** | |
| --- | --- | --- |
|  | F statistic | P-value |
| *My. trossulus* | F_1,4_ = 512.89 | **<0.001** |
| *Ma. gigas* | F_1,4_ = 225.43 | **<0.001** |
| *N. lamellosa* | F_1,4_ = 11.62 | 0.054 |
| *Li. sitkana* | F_1,8_ = 11.85 | **0.035** |
| *Lo. persona* | F_1,4_ = 85.77 | **0.005** |
| *E. viride* | F_1,4_ = 18.49 | **0.040** |
| *B. glandula* | F_1,4_ = 22.84 | **0.026** |
| *H. oregonensis* | F_1,8_ = 6.98 | 0.059 |
| *A. domesticus* | F_1,4_ = 0.03 | 0.863 |

Table S4: Effect of heat denaturation treatment on IRI activity of homogenate in intertidal species. Statistics are from a mixed effects model, with individual ID as a random effect. P-values are reported to three decimal places and have been adjusted using the “Hommel” p-value adjustment.

| **Species** | **Effect of heat treatment on IRI activity** | |
| --- | --- | --- |
|  | F statistic | P-value |
| *My. trossulus* | F_1,4_ = 78.01 | **0.006** |
| *Ma. gigas* | F_1,8_ = 44.45 | **0.001** |
| *N. lamellosa* | F_1,8_ = 4.87 | 0.117 |
| *Li. sitkana* | F_1,3.1085_ = <0.01 | 0.972 |
| *Lo. persona* | F_1,8_ = 83.03 | **<0.001** |
| *E. viride* | F_1,3_ = 34.41 | **0.049** |
| *B. glandula* | F_1,4_ = 6.41 | 0.129 |
| *H. oregonensis* | F_1,4_ = 50.10 | **0.013*** |
| *A. domesticus* | F_1,4_ = 11.95 | 0.086 |

Table S5: Summary of IBP and whole organism results. Columns indicating IBP activity represent if activity could be accounted for by active protein (from Table 2.2 and Table 2.3). SCP column shows supercooling points of whole organisms in °C ± standard error. Final column represents percentage of individuals that survived freezing at -7 °C within 6 days of the cold exposure.

| **Phylum** | **Species** | **INP activity** | **IRI activity** | **Ice shaping** | **SCP (°C)** | **Percent survival after freezing** |
| --- | --- | --- | --- | --- | --- | --- |
| Mollusca | *M. trossulus* | Yes | Yes | Yes | -4.52 ± 0.18 | 100% |
| Mollusca | *M. gigas* | Yes | Yes | Yes | -2.61 ± 0.29 | 100% |
| Mollusca | *N. lamellosa* | Yes | No | No | -5.18 ± 0.29 | 11.11% |
| Mollusca | *Li. sitkana* | Yes | No | No | -5.86 ± 0.12 | 100% |
| Mollusca | *Lo. persona* | Yes | Yes | Yes | -5.18 ± 0.22 | 100% |
| Nemertea | *E. viride* | Yes | No | No | -5.58 ± 0.25 | 50% |
| Arthropoda | *B. glandula* | Yes | No | No | -5.14 ± 0.14 | 83.33% |
| Arthropoda | *H. oregonensis* | No | No | Yes | -5.86 ± 0.21 | 0% |
| Arthropoda | *A. domesticus* | No | No | No | -12.0 ± 0.42 | 0% |

*Notes that 58% of individuals died ten days after freezing.
